# Imaging ATP Consumption in Resting Skeletal Muscle: One Molecule at a Time

**DOI:** 10.1101/2020.05.27.119065

**Authors:** Shane R. Nelson, Amy Li, Samantha Beck-Previs, Guy G. Kennedy, David M. Warshaw

**Author notes:** Department of Pharmacy & Biomedical Sciences, La Trobe University, Bendigo VIC 3550 Australia.

## Abstract

Muscle contraction is driven by sarcomere shortening and powered by cyclic hydrolysis of ATP by myosin molecular motors. However, myosin in relaxed muscle continues to slowly hydrolyze ATP, analogous to an idling engine. Utilizing super-resolution microscopy to directly image single molecule fluorescent ATP turnover in relaxed rat soleus skeletal muscle sarcomeres, we observed two rates of myosin ATP consumption that differed 5-fold. These distinct hydrolysis rates were spatially segregated, with the slower or “super relaxed” rate localized predominantly to the sarcomere C-zone, where Myosin Binding Protein-C (MyBP-C), a known modulator of muscle contractile function, exists. This super relaxed hydrolysis rate and its location suggest that MyBP-C can sequester myosin motors to regulate muscle metabolism and heat production in resting muscle and force generation upon activation.

**One Sentence Summary:** Super relaxed skeletal muscle myosin is stabilized by Myosin-Binding Protein C as imaged by single ATP molecule consumption.

## Main Text

In skeletal muscle’s most basic contractile unit, the sarcomere (Fig. 1A, 1B), force and motion are generated by “active” myosin molecular motors of the thick filament cyclically interacting with actin thin filaments. To power this contractile process, myosin motors hydrolyze adenosine triphosphate (ATP) (Fig. 1E), with approximately one third of the chemical energy converted into mechanical work while the remainder is lost as heat (*1*). Under non-contracting or “relaxed” conditions, myosin is prevented from interacting with actin, due to the presence of calciumdependent regulatory proteins on the thin filament (Fig. 1D). Even when muscle is relaxed, the myosin motors continue to hydrolyze ATP, analogous to the consumption of fuel by an idling car, but at an ATPase rate 2 to 3 orders of magnitude slower than during contraction (*2*). Since skeletal muscle comprises approximately 40% of an adult human’s mass, muscle’s energy expenditure and heat generated while relaxed are major contributors to an individual’s most basic physiological parameters of resting metabolic rate and thermogenesis.

**Figure 1.**
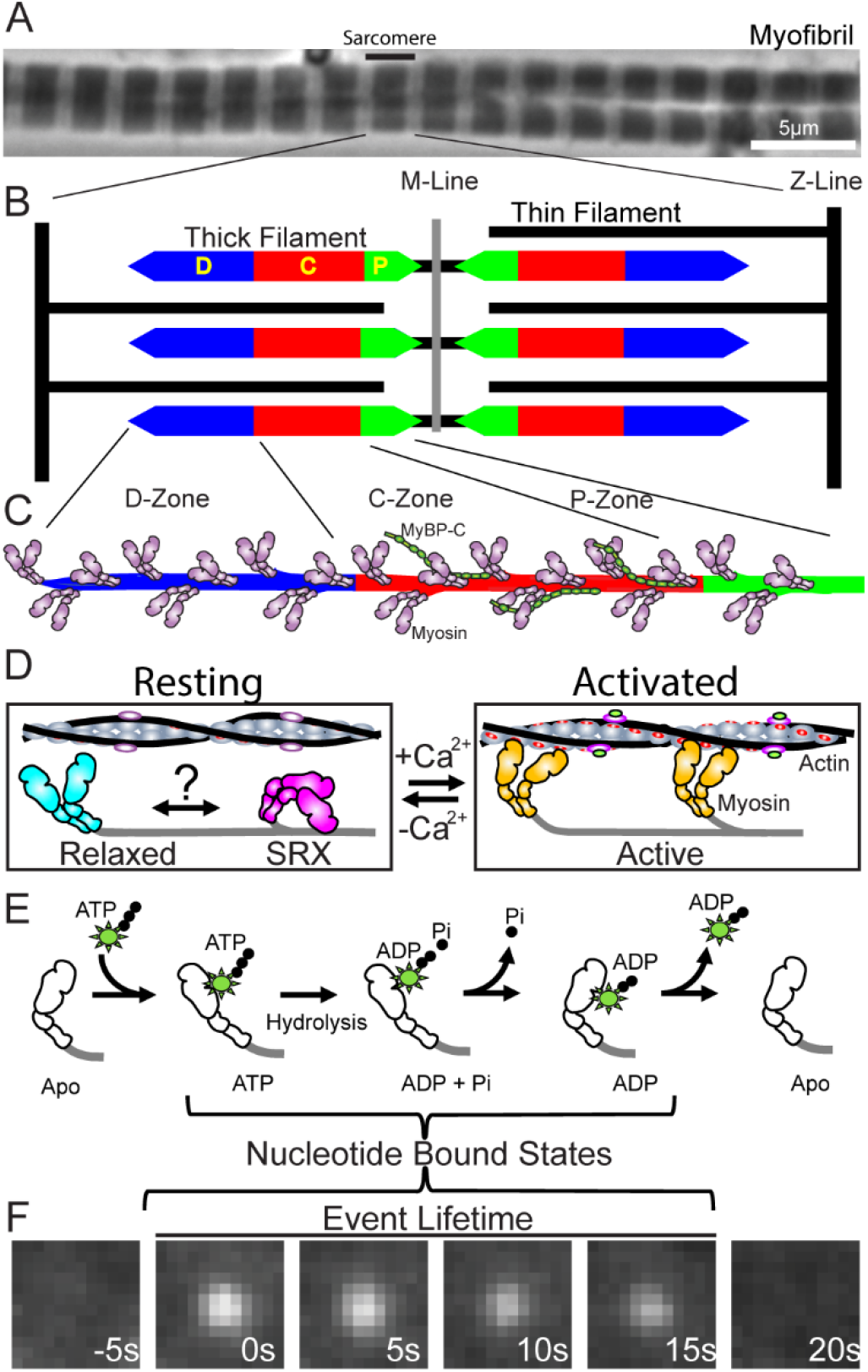
Skeletal muscle myofibrils, sarcomeres and myosin states. **(A)** Phase contrast image of rat soleus myofibrils with sarcomere identified. **(B)** Sarcomere illustration, highlighting interdigitating myosin-based thick filaments and actin-based thin filaments. Thick filaments are bipolar with the M-Line at their center. Each thick filament half is described by a D-, C-, and P-zone indicated by the different colors. **(C)** Illustration of a half-thick filament, with myosin motors protruding from the thick filament backbone and MyBP-C only within the C-zone, where it is found in a 1:3 stoichiometry with myosin. **(D)** In the absence of calcium in the Resting condition (-Ca^2+^), binding sites on thin filament are blocked by troponin-tropomyosin and myosin can adopt either a disordered “Relaxed” or folded “SRX” state that are in a potential (?) dynamic equilibrium. Under Activated conditions in the presence of calcium (+Ca^2+^), binding sites on thin filament are exposed (red), and “Active” myosin is free to bind actin and cause filament sliding. **(E)** Myosin’s ATP hydrolysis cycle. Upon binding a molecule of fluorescently labeled ATP, the immobilized fluorophore can be visualized as shown in **(F)**. The duration of the observed fluorescence spot is a direct readout of the total lifetime of the nucleotide bound states.

Curiously, myosin’s basal ATPase rate in the absence of actin is 5-fold greater *in vitro* than measured in muscle fibers (*3,4*), implying that an additional inhibitory mechanism must exist to lower energy usage within relaxed muscle. Therefore, recent models that are based on muscle fiber ATP consumption, suggest that two myosin populations exist at rest, and have kinetically distinct hydrolysis rates: 1) a relaxed myosin population (Fig. 1D), consistent with the previously described *in vitro* basal rate, and; 2) a significantly inhibited myosin population, termed “super relaxed” or “SRX” myosin (Fig. 1D) (*5*). SRX myosin may represent a “sequestered” or “off” state, which is stabilized by intramolecular interactions that form when myosin molecules are packed into thick filaments within the sarcomere (Fig. 1C, 1D) (*6*). Thus, the SRX state represents a potentially important mechanism for conserving energy in relaxed muscle and for regulating force generation in active muscle through sequestration of myosin motors. Is the SRX state itself regulated? Does this impact both skeletal muscle’s resting metabolic rate and force generated during a contraction? Myosin-binding protein C (MyBP-C), a thick filament-associated protein (Fig. 1C), may serve this regulatory role by stabilizing the myosin SRX state through its molecular interactions with the myosin molecule (*7*).

The importance of MyBP-C as a regulator of skeletal muscle function is emphasized by genetic mutations in MyBP-C leading to distal arthrogryposis, a skeletal muscle myopathy characterized by pathological hyper-contraction (*8,9*). Interestingly, MyBP-C is anchored by its C-terminus to the backbone of the myosin thick filament, but only within a limited region known as the “C-zone” (Fig. 1B, 1C) (*10*). Therefore, does MyBP-C’s spatial confinement suggest that its putative role in sequestrating myosin motors into the SRX state occurs only within the C-zone and not within flanking regions of the thick filament that are devoid of MyBP-C (i.e. D- and P-zones; Fig. 1B, 1C)? If this is not the case, then are all myosin motors within the thick filament capable of adopting the SRX state, regardless of their location? To address these questions, we spatially resolved myosin’s hydrolysis of individual fluorescently-labeled ATP in relaxed skeletal muscle sarcomeres, allowing us to identify the presence of SRX myosin motors and their location within the thick filament.

Myosin’s ATPase rate in the absence of actin is rate-limited by the release of its hydrolysis products, inorganic phosphate (Pi), followed by the subsequent release of adenosine diphosphate (ADP) (Fig. 1E) (*11*). Thus, fluorescently-labeled ATP in demembranated, relaxed skeletal muscle fibers was used previously to monitor myosin ATPase rates, either by measuring the uptake of fluorescent-ATP or the release of the hydrolysis product, fluorescent-ADP (*5*). By this approach, the myosin SRX state was discovered (*5*). However, by using these large muscle fiber preparations, it was impossible to resolve the spatial location of SRX myosin motors within the sarcomere and specifically, if they are enriched in the C-zone where MyBP-C resides. Therefore, we isolated myofibrils from rat soleus, a predominantly slow-twitch muscle. Myofibrils are muscle contractile organelles, consisting of long, repeating arrays of sarcomeres that are clearly visible as alternating light and dark bands using phase contrast microscopy (Fig. 1A).

Myofibrils were adhered to a glass coverslip and exposed to Relaxing Buffer (0mM Ca^2+^, 4mM ATP, see Supplementary Materials) at room temperature. Under these zero calcium conditions, myofibrils were relaxed as myosin was prevented from attaching to actin by the calciumdependent, thin filament regulatory proteins (Fig. 1D). However, as described above, myosin motors in relaxed sarcomeres continue to hydrolyze ATP. Myofibrils were then exposed to a limited amount (10nM) of fluorescent boron-dipyrromethene-labeled ATP (BODIPY-ATP) so that 1 out of every 400,000 ATP molecules were fluorescent. This permitted single molecule imaging of transient fluorescent nucleotide binding within the myofibril (Fig. 1F; Movie S1). In control experiments, BODIPY-ATP was indistinguishable from unlabeled ATP as a substrate for myosin activity in an *in vitro* motility assay (See Supplementary Materials). Since fluorescent nucleotide binding events were only seen in myofibrils where myosin is the predominant ATPase, these fluorescent nucleotide binding events directly report the nucleotide-bound states of myosin’s ATPase cycle (Fig. 1E, 1F). Specifically, appearance of a stationary BODIPY-ATP reflects its binding to myosin’s apo state, while the lifetime of the fluorescence event encompasses hydrolysis, Pi release, and ultimately the release of BODIPY-ADP (Fig. 1E, 1F). As product release is the rate-limiting step of the hydrolytic cycle, the fluorescence event lifetime (Fig. 1F; white streaks in kymograph, Fig. 2A) serves as an estimate of the myosin ATPase cycle time. The fit to the survival plot of fluorescence event lifetimes (n=734 events from 259 sarcomeres from 20 myofibrils) was best described by the sum of two exponential populations (ATP, Fig. 2B), as shown by reduced residuals from the fits (Fig. 2C). The time constants differ by 5-fold (i.e., 26±2s and 146±16s), with the longer lifetime population accounting for 28±4% of the total (inset, Fig. 2B; Table 1). These time constants are in remarkable agreement with fiber-level measurements in rabbit soleus muscle (*5, 12*), suggesting that the longer lifetime population is reflective of SRX myosin in the myofibril preparation.

**Figure 2.**
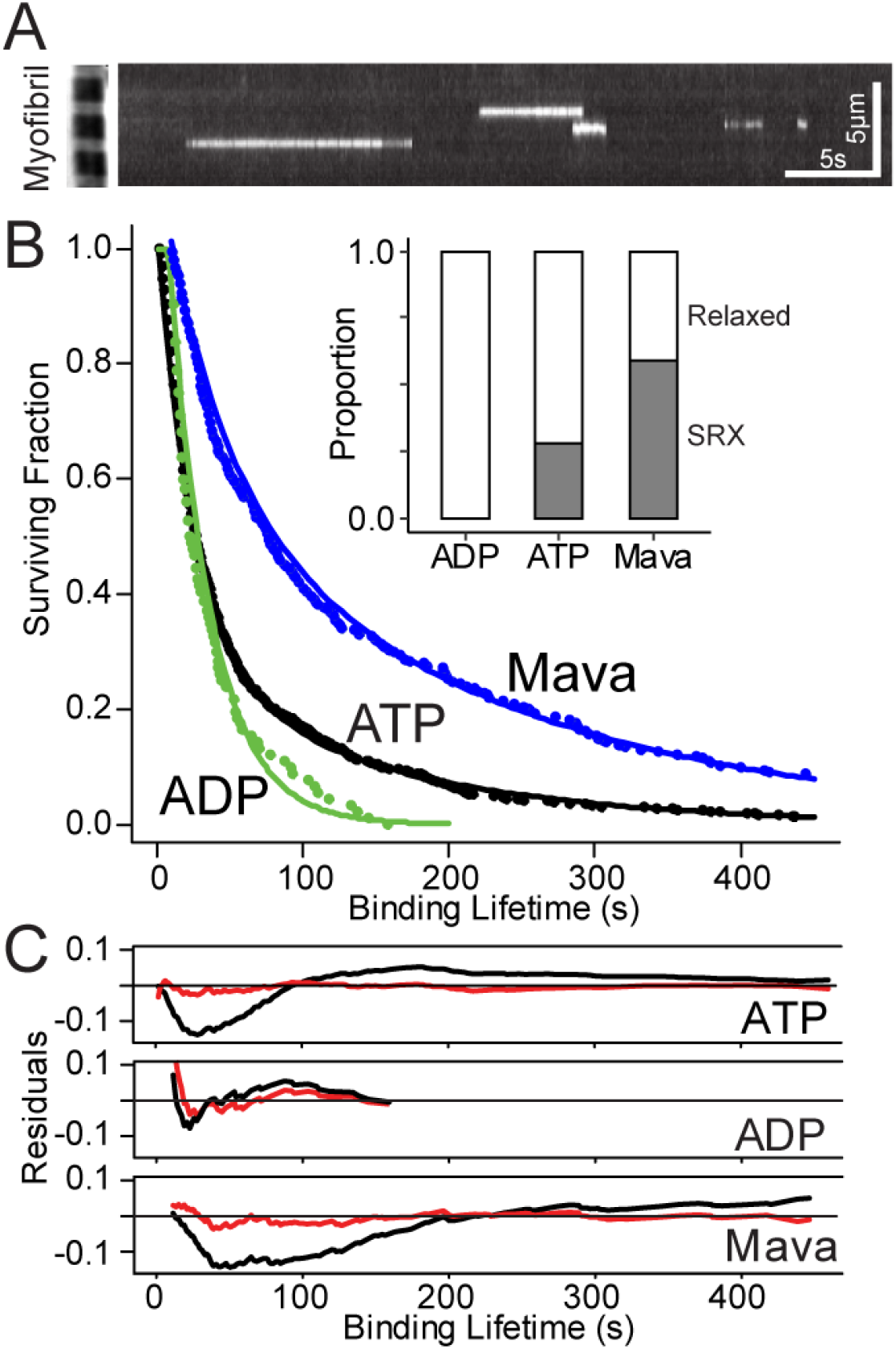
Fluorescent-nucleotide binding lifetimes. (**A**) Kymograph showing multiple fluorescent-nucleotide binding events (individual streaks), with phase contrast image of a myofibril provided (left side) for scale. **(B)** Survival plot of fluorescent-nucleotide binding lifetimes for myofibrils incubated in ADP (green), ATP (black), or ATP+Mavacamten (Mava) (blue). Lines demonstrate double exponential fits to each dataset and inset shows relative proportions of relaxed (white) and SRX (grey) events for each experimental condition. **(C)** Residuals from fitting each experimental condition are reduced and lack clear structure when fit with double (red) versus single (black) exponential.

**Table 1.**
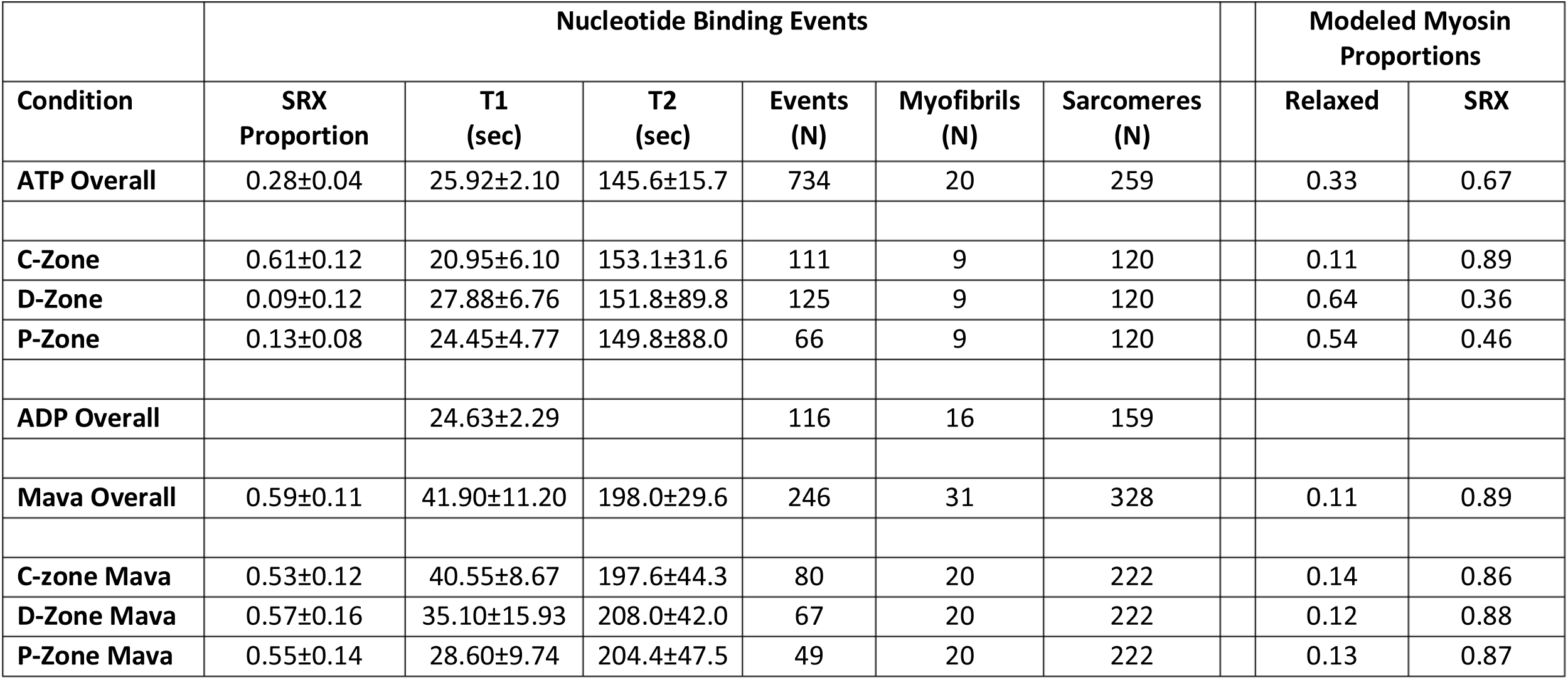
Fluorescent-Nucleotide Binding Events and Modeled Myosin Proportions. “SRX Proportion” is the proportion of events attributable to the SRX population, “Tl” and “T2” are time constants for the “relaxed” and SRX populations, respectively, from fitting event lifetime data with double exponential model, ± values are estimated standard errors of the fit (see Supplementary Materials), N is the number of total events, myofibrils or sarcomeres. Modeled Myosin Proportions are calculated based upon the “Myosin State Proportions Model,” details in Supplementary Materials. “Mava” denotes myofibrils treated with Mavacamten.

| Nucleotide Binding Events |  |  |  |  |  |  | Modeled Myosin Proportions |  |
| --- | --- | --- | --- | --- | --- | --- | --- | --- |
| Condition | SRX Proportion | T1 (sec) | T2 (sec) | Events (N) | Myofibrils (N) | Sarcomeres (N) | Relaxed | SRX |
| ATP Overall | 0.28±0.04 | 25.92±2.10 | 145.6±15.7 | 734 | 20 | 259 | 0.33 | 0.67 |
| C-Zone | 0.61±0.12 | 20.95±6.10 | 153.1±31.6 | 111 | 9 | 120 | 0.11 | 0.89 |
| D-Zone | 0.09±0.12 | 27.88±6.76 | 151.8±89.8 | 125 | 9 | 120 | 0.64 | 0.36 |
| P-Zone | 0.13±0.08 | 24.45±4.77 | 149.8±88.0 | 66 | 9 | 120 | 0.54 | 0.46 |
| ADP Overall |  | 24.63±2.29 |  | 116 | 16 | 159 |  |  |
| Mava Overall | 0.59±0.11 | 41.90±11.20 | 198.0±29.6 | 246 | 31 | 328 | 0.11 | 0.89 |
| C-zone Mava | 0.53±0.12 | 40.55±8.67 | 197.6±44.3 | 80 | 20 | 222 | 0.14 | 0.86 |
| D-Zone Mava | 0.57±0.16 | 35.10±15.93 | 208.0±42.0 | 67 | 20 | 222 | 0.12 | 0.88 |
| P-Zone Mava | 0.55±0.14 | 28.60±9.74 | 204.4±47.5 | 49 | 20 | 222 | 0.13 | 0.87 |

Upon muscle activation as cytoplasmic calcium rises, SRX myosin must rapidly transition to the “relaxed” state and subsequently to the “active” state (Fig. 1D) in order to generate force and motion (*13*). In fact, in myofibril experiments where the 1mM unlabeled ATP was replaced with 1mM unlabeled ADP so that myosin strong-binding was favored (*14*), the myofibril was unable to relax and thus, myosin was prevented from entering the SRX state (*5*). When these myofibrils were exposed to 10nM fluorescent BODIPY-ATP, myofibrils slowly contracted at 0.002 sarcomere lengths/s. Under these non-relaxing conditions, the fluorescence event lifetimes (n=116 events from 159 sarcomeres from 6 myofibrils) were best fit by a single population with lifetimes (i.e. 25±2s) equal to the shorter lifetimes measured in the relaxed myofibrils, with no evidence of an SRX myosin population (ADP, Fig. 2B; Table 1). Most likely some myosin motors hydrolyzed BODIPY-ATP at the “active” actin-activated rate. However, this rate would be faster than the camera’s 1s^-1^ framerate and thus undetected (*1, 13*). Regardless, the observed kinetics suggest that the long-lived fluorescence lifetimes are sensitive to the activation state of the myofibril, as predicted for the SRX state, and not merely an artifact of non-specific BODIPY-ATP binding. Further evidence to support the myosin-specific nature of these events was with the use of Mavacamten; a myosin-selective, small molecule ATPase inhibitor (*15*). Mavacamten slows the rate of both Pi and ADP release in both cardiac and skeletal muscle (*11*) by stabilizing the myosin SRX state (*16*). When resting myofibrils were treated with 10μM Mavacamten, once again the survival plot of the fluorescence lifetimes (n=246 events from 328 sarcomeres from 31 myofibrils) were best fit by two populations with time constants approximately 50% longer than in the absence of Mavacamten (Mava, Fig 2B). Additionally, the overall proportion of SRX events doubled from 28±4% to 59±11% (Inset, Fig. 2B; Table 1), confirming that the SRX population originates from myosin and that Mavacamten does indeed enhance the presence of this inhibited myosin state.

We then took advantage of the myofibril’s well-defined structure to spatially localize where SRX myosin exist within the sarcomere. More specifically, if MyBP-C can sequester myosin into the SRX state, do these myosins exist exclusively within the C-zone where MyBP-C is found (Fig. 1C) and if not are SRX myosin distributed along the entire length of the thick filament? Sarcomeres exhibit bilateral, structural symmetry about the M-line at the center of the sarcomere (Fig. 1B), where myomesin, an Ig-superfamily protein, exists and serves to connect adjacent thick filaments (*17*). By identifying sarcomere centers through myomesin immunofluorescence (Fig. 3A), the center-to-center distance averaged 2.2±0.2μm, consistent with relaxed sarcomere lengths in skeletal muscle (*18*). Then fluorescent BODIPY-ATP binding events and myomesin (M-line) were localized with high spatial resolution (Fig. 3A) using super-resolution detection algorithms (see Supplementary Materials). After color registration and drift correction (see Supplementary Materials), we measured the distance between each BODIPY-ATP binding event and the M-line with a combined error (σ) of 38nm. Due to the sarcomere’s structural symmetry, we could combine events from both sides of the M-line with those from multiple sarcomeres (n=120) and multiple myofibrils (n=9). Once combined (n=302 events), fluorescence binding events appeared randomly distributed up to 971nm from the M-line (Fig. 3B). With the distance from the M-line to the thick filament end being 817nm (*19*) and allowing for localization error (σ), 97% of the localized BODIPY-ATP binding events lie within the span of the myosin thick filament, providing further evidence for the myosin-specific nature of these events.

**Figure 3.**
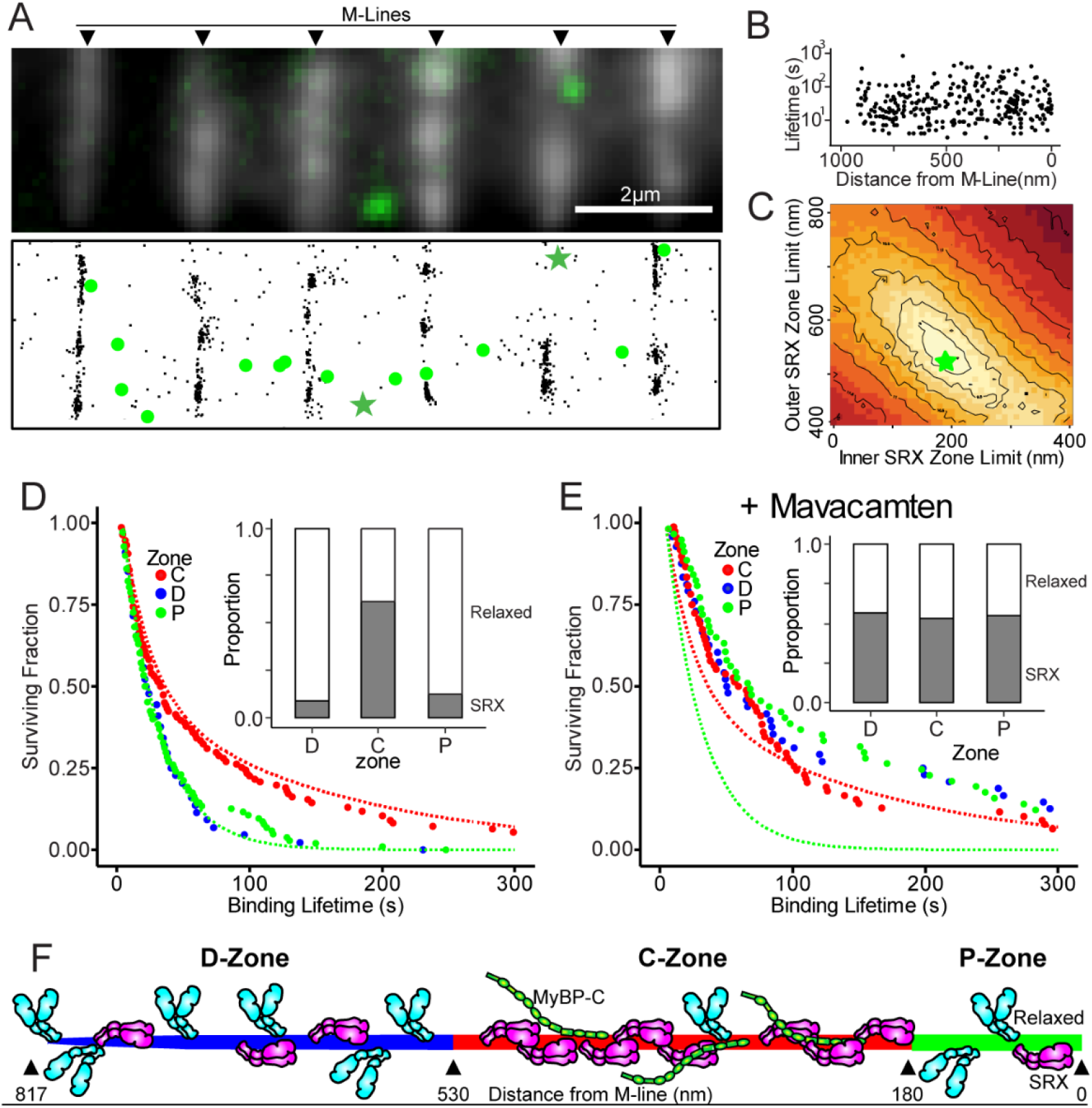
Spatial localization of fluorescent nucleotide binding events. **(A)** Top panel: fluorescence image of anti-myomesin labeled (M-line) myofibrils with overlaid still frame showing two fluorescent-nucleotide binding events (larger green spots). Bottom panel: color- and drift-corrected sub-pixel localizations for myomesin (black points) and fluorescent-nucleotide events (green), with binding events from upper panel denoted (green stars). **(B)** Scatterplot of fluorescent-nucleotide binding event lifetimes vs. distance from nearest M-line. **(C)** RMSD heatmap comparing a simulated SRX-containing zone of variable width (i.e. bounded by inner and outer limits) to experimental data (see Supplementary Materials). Lighter colors denote better fits, with best fit (green star, SRX Zone Limits: Inner: 180nm, Outer: 530nm). **(D)** Survival plots of binding event lifetimes localized to D-, C- and P-zones. Dashed lines indicate theoretical curves if a zone contained only relaxed events (green line) or entirety of the SRX events (red line). Inset bar chart shows proportions of Relaxed (white) and SRX (grey) events by zone. **(E)** As in (D), but for myofibrils incubated in ATP with Mavacamten. **(F)** Cartoon illustrating that C-zone contains the preponderance of sequestered SRX myosin (magenta, folded), while D- and P-zones contain predominantly relaxed myosin (cyan, extended). Distances for zones determined by modeling effort in (C).

Although no spatial bias was apparent for the BODIPY-ATP binding events (Fig. 3B), could the long-lived SRX event population be localized to a specific region of the sarcomere and thus the thick filament? To address this, we took an unbiased approach and developed a simple “Monte Carlo-style” model that simulated fluorescent nucleotide binding events along the length of a half thick filament. These events were assumed to originate from two populations having relative proportions and time constants observed experimentally (see model details in Supplementary Materials). We then compared the experimental data to simulated results over a wide range of possible bounds for an SRX-containing zone. From this comparison a landscape emerges where the best fit to the experimental data suggests that SRX events occurred between 180-530nm from the M-line (Fig. 3C). This predicted 350nm SRX-containing zone is in remarkable agreement with the location of the C-zone that we recently reported in rat soleus fibers, based on MyBP-C immunofluorescence (*19*) and from previous immuno-electron microscopy of rabbit psoas muscle (*10*). Using this model prediction, we parsed the experimentally observed nucleotide binding events into the C-zone (180-530nm from the M-line, n=111) and its flanking D-zone (> 530nm from the M-line, n=125) and P-zone (< 180nm from the M-line, n=66), which are both devoid of MyBP-C (Fig. 3F). We then fit the survival plots of the fluorescence lifetimes for each of these three zones to double exponentials (Fig. 3D). Based on these fits, the C-zone was characterized as having 61±12% SRX events, while the flanking D- and P-zones were characterized by 9±12% and 13±8% SRX, respectively (Fig 3D; Table 1). Thus, the SRX state appeared to be enriched within the C-zone.

Prior muscle fiber studies using fluorescent MANT-ATP to identify the presence of the SRX state were performed as single-turnover experiments (*5*). In these studies, the observed proportions of the fast and slow components of the fluorescent MANT-ATP decay following an unlabeled ATP chase directly reflected the underlying myosin populations and their hydrolytic rates. However, in our myofibril assay, all myosins are constantly turning over ATP and only occasionally hydrolyze a fluorescent BODIPY-ATP molecule. Therefore, the slower SRX myosin population will cycle through approximately 5-fold less ATP in the same time as the faster “relaxed” myosin population. Thus, the proportion of observed SRX-related fluorescence lifetime events, based on our double exponential fits (Figs. 2B, 3D, 3E), underestimates the underlying SRX myosin population. Therefore, we developed a simple model (see details in Supplementary Materials) to account for this discrepancy, wherein the observed frequency of fluorescent nucleotide binding events is a function of both the underlying proportion of the myosin populations (relaxed or SRX), as well as the ratio of the hydrolytic rates between the two myosin populations. Using this model, we estimate that 89% of the myosin within the C-zone are in the SRX state, compared to ~40% within each of the D- and P-zones (Table 1) as illustrated in Figure 3F. This prediction suggests that every myosin motor along the length of the thick filament may be in a dynamic equilibrium between a relaxed and SRX state (Fig. 1D) but that this equilibrium is shifted towards the SRX state within the C-zone where MyBP-C only resides. Interestingly, within the C-zone there is only 1 MyBP-C molecule for every 3 myosin molecules (*13*). If MyBP-C is responsible for the large proportion of SRX myosin within the C-zone, then each molecule of MyBP-C may stabilize more than one myosin molecule into the SRX state through a cooperative mechanism, such as the C terminus of MyBP-C interacting with the tails of multiple myosins in the thick filament. Finally, across the entire sarcomere, we estimate that 68% of myosin is in the SRX state. This estimate is higher than previously reported values of 35-50% from skinned fiber studies (*5*), however, closer to the estimated value needed to match the low ATP consumption from intact relaxed muscle (*13*).

When myofibrils were treated with 10μM Mavacamten, the proportion of SRX events doubled (Fig. 2B; Table 1). So where along the thick filament length did the increased SRX population originate? When BODIPY-ATP binding events (n=246 from 222 sarcomeres and 20 myofibrils) were spatially mapped by their distance from the M-line, the proportion of SRX events within the C-zone (53±12%) was not significantly different compared to untreated myofibrils (p> 0.05, Table 1). However, the events within the D- and P-zones demonstrated a marked shift towards the SRX state (Table 1; Fig. 3D, 3E). In fact, when the ATPase turnover model (see above) was used to calculate the SRX myosin population for the various zones, the D- and P-zones estimates of 88% and 87%, respectively, were nearly equal to the 86% predicted for the C-zone (Table 1), which is visually discernable by comparing the fluorescence lifetime plots in Figure 3D versus Figure 3E. Thus, the increased SRX myosin population upon treatment with Mavacamten was restricted to the regions flanking the C-zone, further confirming that myosin motors along the entire length of the thick filament are capable of entering the SRX state, regardless of whether or not MyBP-C is present. However, without pharmacologic intervention, the SRX myosin state is enriched only within the C-zone, strongly supporting MyBP-C’s role in stabilizing the SRX state.

If in relaxed muscle a significant proportion of myosin motors are sequestered into the SRX state, then by what mechanism do these motors transition into states that can generate force upon muscle activation? Historically, activation of skeletal muscle was thought to be thin filament-based, whereby calcium released from the sarcoplasmic reticulum binds to the thin filament regulatory protein complex (troponin-tropomyosin), causing a shift in the position of this complex so that myosin motors can bind to the thin filament and generate force (Fig. 1D) (*20, 21*). This activation scheme assumes that myosin motors are always “on” and available to bind to the thin filament. However, recent models suggest that muscle activation may involve a thick filament-dependent mechanism as well (*22*). Specifically, strain in the thick filament may be the “mechanosensor” that serves as the trigger to release myosin from the SRX or “off” state, allowing these myosins to contribute to force generation (*22*). Interestingly, thin filament activation in skeletal muscle may spatially begin near the D-zone, as this is the location of calcium release from the sarcoplasmic reticulum (*23*). If so, then myosin motors within the D-zone, which are predominantly in the relaxed or “on” state (Fig. 3F), would be the first to bind the thin filament. In fact, a recent study in cardiac muscle (*24*) suggests that activation along the thick filament does begin in the D-zone, where myosins that are “constitutively ON” bind to the thin filament and generate force, in turn, producing strain along the entire length of the thick filament (*25*). This strain would facilitate the release of SRX myosin motors, particularly those that are enriched in the C-zone nearer the center of the thick filament. This concept is supported by the dramatic elimination of SRX myosin in myofibrils that are effectively activated in the presence of ADP (Fig. 2B; Table 1).

With the majority of SRX myosin motors being spatially segregated to the C-zone in resting myofibrils, what are the functional implications of this spatial segregation? In skeletal muscle, MyBP-C is believed to be a critical modulator of contractility, serving to simultaneously sensitize the thin filament to calcium through its N-terminal interaction with the thin filament (*19*) and to act as a “brake” to reduce sarcomere shortening (*26*) and thin filament sliding velocity *in vitro* (*27, 19*). Both of these modulatory capacities are observed only within the C-zone where MyBP-C is found (*19*) and may be impacted by the extent to which MyBP-C sequesters myosin into the SRX state. Interestingly, in slow-twitch skeletal muscle, multiple isoforms of MyBP-C are expressed as a result of alternative exon splicing, and we have recently shown that these variants have different capacities to modulate myosin’s motion generating interactions with the thin filament (*19*). These MyBP-C isoforms may also differentially stabilize the SRX state with implications for both relaxed and activated muscle physiology. The impact of MyBP-C on the SRX state is supported by mutations in the cardiac isoform of MyBP-C resulting in reduced proportions of SRX myosin, which may explain the hyper-contractility and impaired relaxation that are characteristic of these cardiomyopathies (*28, 29*). Similarly, patients with genetic mutations in skeletal MyBP-C exhibit distal arthrogryposis, a condition with extreme muscle contractures (*8, 9*), that is potentially caused by the mutant MyBP-C’s inability to sequester myosin motors into the SRX state. In conclusion, the majority of myosin motors in relaxed slow-twitch skeletal muscle are in the SRX state. Therefore, slight shifts in this SRX myosin population may potentially impact muscle’s normal resting metabolism, heat generation, and contractility. Since MyBP-C may act as a spatially localized modulator of the SRX state within the C-zone, where the SRX state is enriched, focus on MyBP-C as a therapeutic target to modulate muscle metabolism and force generation in cases of muscle weakness in the elderly (*30*) or pathologic hyper-contractility (*8,9*) is warranted.

## Acknowledgments

The authors wish to acknowledge George Osol and Marilynn Cipolla (Univ. of Vermont) for contribution of biological samples.

## Authors contributions: Shane Nelson

Methodology, Software, Formal analysis, Investigation, Resources, Writing – original draft, Writing – review and editing, Visualization;

## Amy Li

Investigation, Resources;

## Samantha Beck-Previs

Investigation, Resources;

## Guy Kennedy

Methodology, Resources, Visualization;

## David Warshaw

Conceptualization, Writing – review and editing, Supervision, Project administration, Funding acquisition.

## Data and materials availability

All data are available in the manuscript or the supplementary materials. Software available upon request.

## Supplementary Materials

Materials and Methods

Figures S1-S4

Movie S1

