## Supplementary Materials for "Imaging ATP Consumption in Resting Skeletal Muscle: One Molecule at a Time"

**This PDF file includes:**

Materials and Methods

Figs. S1 to S4

Movie S1

### Materials and Methods

#### Myofibril Preparation

Glycerinated muscle strips were prepared from soleus muscle from female, non-pregnant Sprague Dawley rats. Muscle strips of approximately 1-2mm diameter and 1.5cm in length were tied to toothpicks using silk sutures and washed in Rigor Buffer (50 mM Tris, 100 mM $Nacl_2$, 2mM KCl, 2mM $MgCl_2$, 10 mM EGTA, 0.5mM DTT, protease inhibitor cocktail, pH 7.4) twice for 12 hours each time, at 4°C. This was followed by two washes (12 hours each) with a 50:50 glycerol: Rigor Buffer, and finally stored at -20°C for a minimum of 7 days in 50:50 glycerol: Rigor Buffer.

Myofibrils were prepared from glycerinated muscle strips immediately prior to imaging by dissecting a thin strip (~1mm diameter) and de-glycerinating by soaking in 1.5ml of Myofibril Preparation Buffer (50 mM Tris, 100 mM KCl, 4mM $MgCl_2$, 2mM EGTA, protease inhibitor cocktail, pH 7.0) on ice for 1 hour. Samples were then mechanically sheared using a "Tissue Tearor" homogenizer (Biospec Products, Bartlesville, OK) according to the following speed ramp (in 1000X RPM): 3-9-15-21-15-9-3, for 5 seconds at each speed. Samples were then allowed to settle on ice for 10-15 minutes.

#### Imaging Conditions

From the tube of prepared myofibrils, 200µl were drawn from the bottom of the tube and incubated with anti-myomesin antibody (Cat #EPR17322-9, Abcam, Cambridge UK) at a 1:5000 final dilution for 15 minutes on ice, followed by Alexa 647 goat-anti-mouse IgG (Cat #A21237, Life Technologies, Carlsbad CA) at a final concentration of 1:5000. After 5 minutes on ice, labeled myofibrils were flowed into flowcells, prepared as described previously *(31)*, which were made using plasma-cleaned glass. The flowcell was then incubated at room temperature for 20 minutes to allow myofibrils to adhere to the coverglass surface. Finally, 100µl of Relaxing Buffer (120mM KOAc, 5mM K-phosphate, 4mM EGTA, 4mM $MgCl_2$, 50mM MOPS, 4mM ATP, 10nM BODIPY-ATP (Cat #A12410, ThermoFisher, Waltham MA), pH 6.8) was flowed into the flowcell and samples were imaged using a custom-built dual camera super-resolution microscope, described previously *(32)* for 50 minutes at 10FPS. From each myofibril preparation 2-4 recordings were made, with each flowcell being used for only a single imaging session.

#### Image Analysis

BODIPY-ATP binding lifetimes and positions were measured separately to maximize accuracy and precision. Lifetimes were measured manually from kymographs to account for visually apparent blinking behavior, while position information was calculated using ThunderSTORM localizations, along with drift and channel offset corrections. Details below:

BODIPY-ATP binding events (green channel) were manually documented in image stacks using ImageJ *(33)*. First, image stacks were integrated to 1FPS using the "grouped Z-project" function. Image stacks were then converted into a stack of kymographs using the "Reslice" function and events were documented using rectangular selections and the "overlay" function.

Sub-pixel localizations of both BODIPY-ATP and Myomesin label (red channel) were performed using ThunderSTORM plugin *(34).* Finally, BODIPY-ATP binding lifetimes (from kymograph analysis) were connected to sub-pixel localizations, as the kymograph X, Y, and Slice coordinates can be directly related to Timestamp, X, and Y coordinates (respectively) in the ThunderSTORM output files.

Positions of each M-line were calculated by regression of a line segment by adjusting the segment's defining coordinates ($X_1$, $Y_1$, $X_2$, $Y_2$) to minimize the summed distance squared between each ThunderSTORM localization and the segment. Red channel localizations from only the initial 200 seconds of the recording were used, due to photobleaching, as well as to minimize the blurring effects of stage drift

Drift correction was performed by tracking a fiducial marker (any persistent spot in either red or green channel) over the timecourse of the experiment, followed by linear (occasionally piece-wise) regression to X vs Time and Y vs Time data separately. Coordinates for BODIPY-ATP localizations were then drift corrected by subtracting the calculated drift that occurred between the start of the recording and the onset of each BODIPY-ATP binding event.

Spatial offsets between red and green imaging channels were calculated by imaging multicolor-fluorescent beads (Tetraspeck, ThermoFisher) and analysis using the Particle Image Velocimetry plugin for ImageJ *(35),* which calculates relative offsets between the two images by maximizing cross-correlation between corresponding regions in the two images.

Finally, coordinates of BODIPY-ATP binding events (green channel) were drift- and color-corrected to pixel coordinates of the Myomesin (red channel) at the start of the recording, and distances were calculated between each binding event and the fitted M-line segment.

Error propagation of the combined error for BODIPY-ATP localization, M-line localization, drift, and channel correction yields an overall estimate of 38nm (S.D.) on measured sub-sarcomeric localizations.

**Fluorescent Event Lifetime Fitting**

Events < 3s in duration are rarely detected above background in our analysis, therefore, fluorescence event lifetimes were fit with single and double exponential models that account for a lack of the shortest event lifetimes in our experimental distributions:

Single Exponential:

$$f(x;t) = e^{-x/t} / \left(1 - \left(1 - e^{-\min(x)/t}\right)\right)$$

Double Exponential:

$$f(x;P1,t_1,t_2) = \frac{P_1 e^{-\frac{x}{t_1}}}{\left(1 - \left(1 - e^{-\frac{\min(x)}{t_1}}\right)\right)} + \frac{(1-P_1)e^{-x/t_2}}{\left(1 - \left(1 - e^{-\frac{\min(x)}{t_2}}\right)\right)}$$

Where $P_1$ is the proportion of "relaxed" events, $t_1$ is the time constant for the relaxed population, and $t_2$ is the time constant of the SRX population. Lifetimes were fit by direct optimization of the log-likelihood estimates as implemented in the "fitdistr" function included in the "MASS" library for the statistical programming language "R" (*36*) using the "L-BFGS-B" method *(37).*

**Actin Filament Motility to Validate BODIPY-ATP**

*In vitro* motility was performed using myosin and actin purified from Chicken pectoralis muscle *(38).* Actin was labeled with Alexa 647-phalloidin. Motility experiments were performed as described previously *(31)* at 25, 50, and 100μM ATP or BODIPY-ATP. Movies were recorded at 15FPS on an inverted Nikon DE2000 scope. Three movies were recorded and analyzed per condition, with tracking performed with the "MTrack2" plugin for ImageJ. Velocity vs [ATP] data were fit with a Michaelis-Menten relationship using "Dose-Response Model Fitting" as implemented in the DRM package of the statistical programming language "R" *(36).* Results shown in Figure S1, demonstrating that BODIPY-ATP sustains myosin activity in a nearly identical manner to ATP, even at very limiting [ATP].

**Monte Carlo Simulation**

We developed a model to help determine the spatial arrangement of Relaxed and SRX myosin along a half-thick filament without pre-supposing that these myosin states exist within defined P-, C- and D-zone boundaries. Therefore, we compared the experimental nucleotide binding event localizations (Main text, Fig. 3B) with simulated data sets having alternate spatial arrangements of SRX and Relaxed events within the sarcomere. This enabled us to determine whether the experimental data were in better agreement with alternate spatial localization hypotheses (e.g., uniformly distributed, or in other spatially distinct regions along the thick filament). Therefore, we simulated plots of Fluorescence Binding Lifetime vs Distance that would arise if the SRX events occur within a select region of the half-thick filament. Simulations and comparisons to experimental results were performed as follows:

**1)** Distances of simulated fluorescent nucleotide binding events from the M-line are generated from a uniform distribution between 0 and 817nm (i.e., the half-thick filament length).

**2)** Individual fluorescent nucleotide binding events (n=300 events), along the entire length of the half-thick filament, are initially populated with "Relaxed" myosin state lifetimes by drawing random numbers from an exponential distribution with a time constant (reciprocal rate) of 26s.

**3)** The lifetimes of randomly selected events within a select region, bounded by "Inner" and "Outer" limits (Fig. S2A) were replaced with SRX myosin state lifetimes drawn from an exponential distribution with a time constant of 146s. The number of events that were "converted" to SRX events was fixed at 28% of the total number of events, to match the experimentally observed overall SRX event frequency (Table 1; Main text Fig. 2B).

**4)** The simulated data were compared to the experimental data as follows:

**A)** Each fluorescence binding event has a measured distance from the M-line, measured in nanometers (termed *M*). The entire event population is then split into "proximal" and "distal" sub-populations by a dividing line (*D*, also measured in nm from the M-line).

**B)** Each sub-population of events (defined by $M<D$ and $M>D$) is independently fit with the double exponential model (described above).

**C)** A Ratio of SRX proportions (*R*, as a function of the position of dividing line, *D*) was calculated based on the SRX population in each of two sub-populations according to:

$$R(D) = \frac{P_{SRX,M<D}}{P_{SRX,M>D}}$$

where $P_{SRX}$ is the proportion of SRX events (1-$P_1$ from the double exponential fit to the fluorescence binding lifetime, see above), over the range of $200<D<650$nm. A plot of R(D) for both simulated (red) and experimental data (black) is shown in Fig. S2B

**D)** The difference in R(D) profiles for simulated and experimental data were then calculated as the Root Mean Squared Difference (RMSD)

**5)** The "Inner" bound (from Step 3) was varied in 10nm increments from 0-400nm from the M-line, and the "Outer" bound was varied from 410-810nm in 10nm increments. This allowed for testing of all hypothetical SRX containing zones, up to and including that where SRX events are uniformly distributed across the entire half-thick filament (i.e., bounds of 0 and 810nm, respectively). The RMSD between simulation and experiment as a function of the inner and outer bounds is shown in main text Fig. 3C.

**Myosin State Proportions Model**

Myofibril experiments were all performed at saturating 4mM ATP. Under these conditions, ATP binding is rapid, such that myosin spends the majority of the time in a nucleotide-bound state. In skinned fiber experiments, *(5)*, 96% of myosin are estimated to be bound with ATP under similar conditions reported here. Therefore, the total ATPase cycle time for both "relaxed" and super-relaxed (SRX) myosin should be reflected in our experimentally measured nucleotide-bound lifetimes for these populations. Therefore, during experiments, relaxed myosin molecules should turnover ~5-fold more ATP than SRX myosin. Therefore, we propose a simple model, where the observed "relaxed" event frequency ($P_{1,OBS}$) is related to the proportion of relaxed myosin in the myofibril ($P_1$) and the fluorescence lifetimes of both relaxed and SRX myosin ($T_1$ and $T_2$, respectively):

$$P_{1,OBS} = \frac{P_1/T_1}{(P_1/T_1) + \big((1-P_1)/T_2\big)}$$

The relationship between observed binding events and the underlying myosin proportions are shown in Figure S3.

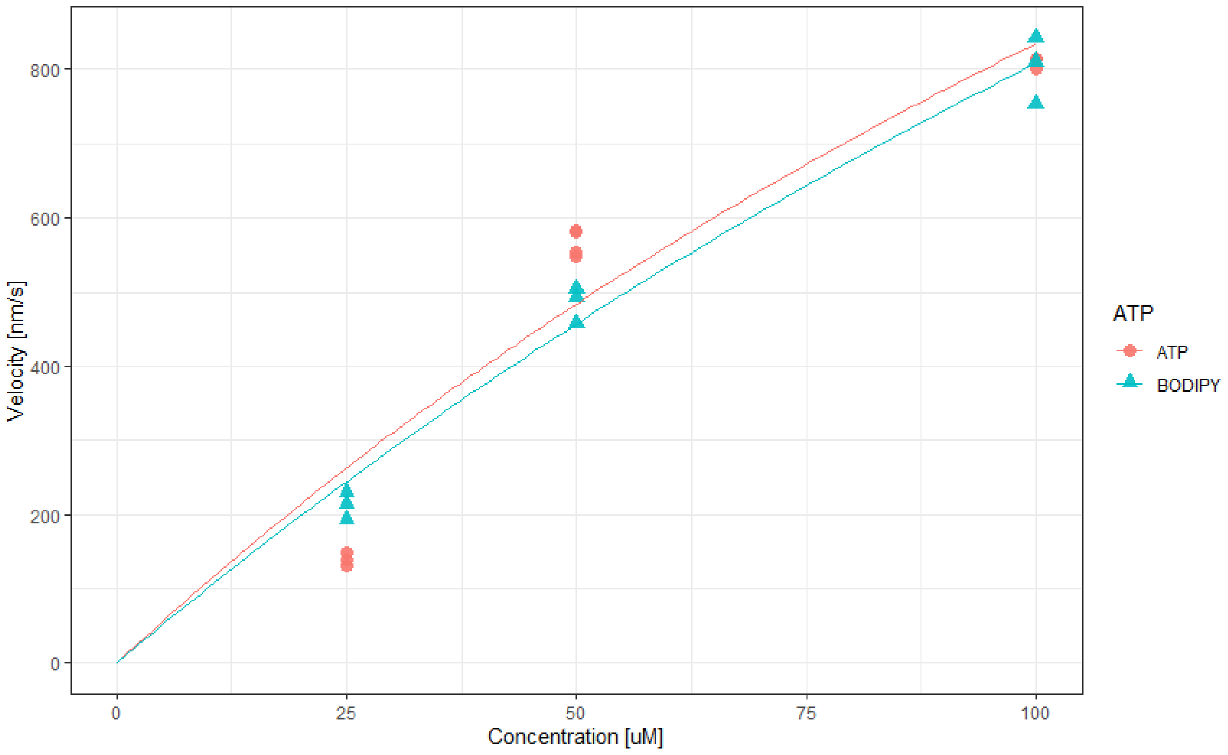


**Figure S1.** Velocity in gliding filament motility assay as a function of concentration of unlabeled ATP and fluorescent BODIPY-ATP. Fitted Values are: $V_{max}$: 3.03±1.82µm/s and $K_m$: 264±204µM for unlabeled ATP and $V_{max}$: 3.49±1.24µm/s and $K_m$: 332±147µM for BODIPY-ATP.

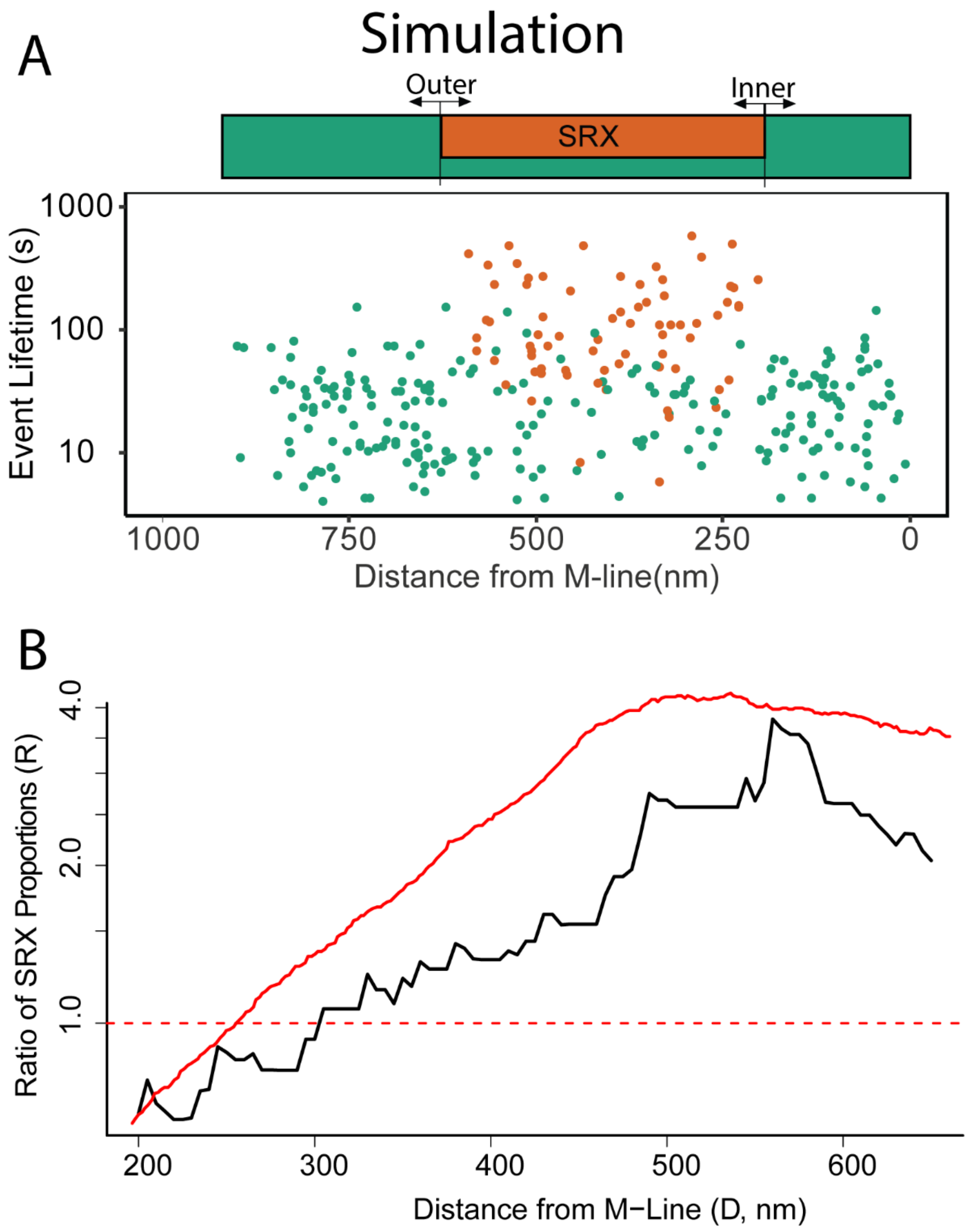


**Figure S2.** Simulation of spatially mapped BODIPY-ATP binding events. (A) Fluorescent-nucleotide binding events were assumed to be uniformly distributed along the length of the half-thick filament (green rectangle). The lifetime for each event (relaxed, green solid circles; SRX, orange solid circles) was determined as described above in Steps 2 and 3. Simulated SRX events occur only within a select region of the half-thick filament (orange box) and are always selected to represent 28% of the total events, in accordance with the experimental data. (B) Profile of R(D) for simulated (red) and experimental data (black).

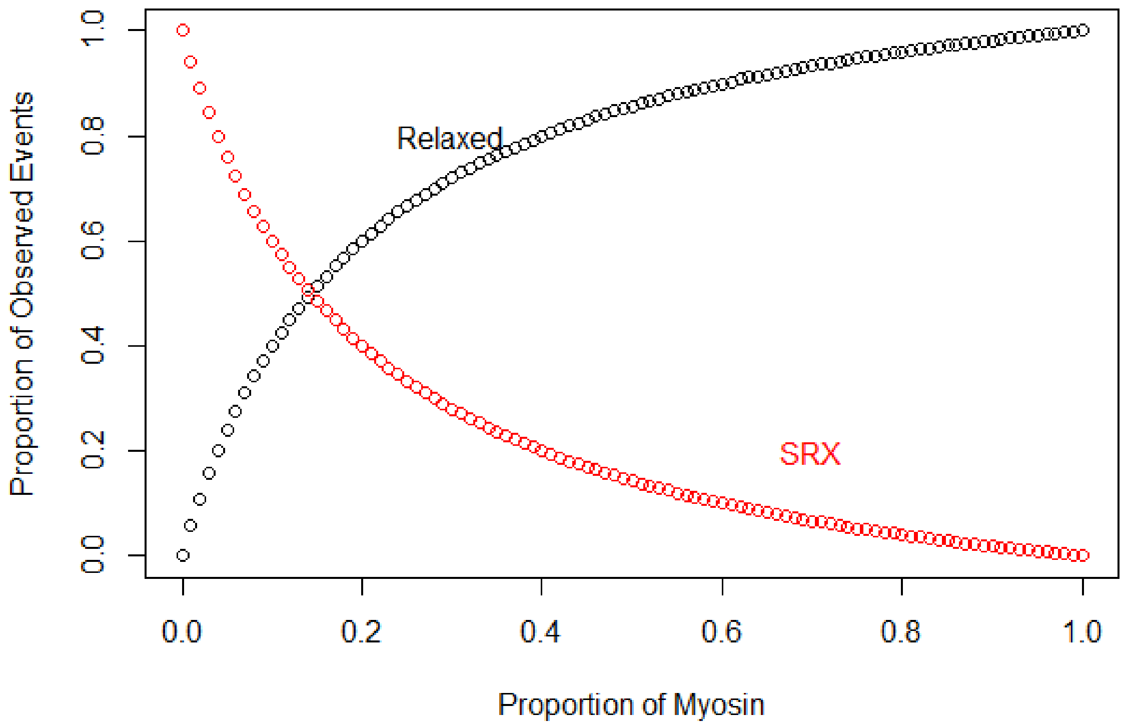


**Figure S3.** Predicted Relaxed/SRX proportion of observed fluorescent BODIPY-ATP binding events as a function of underlying myosin population.

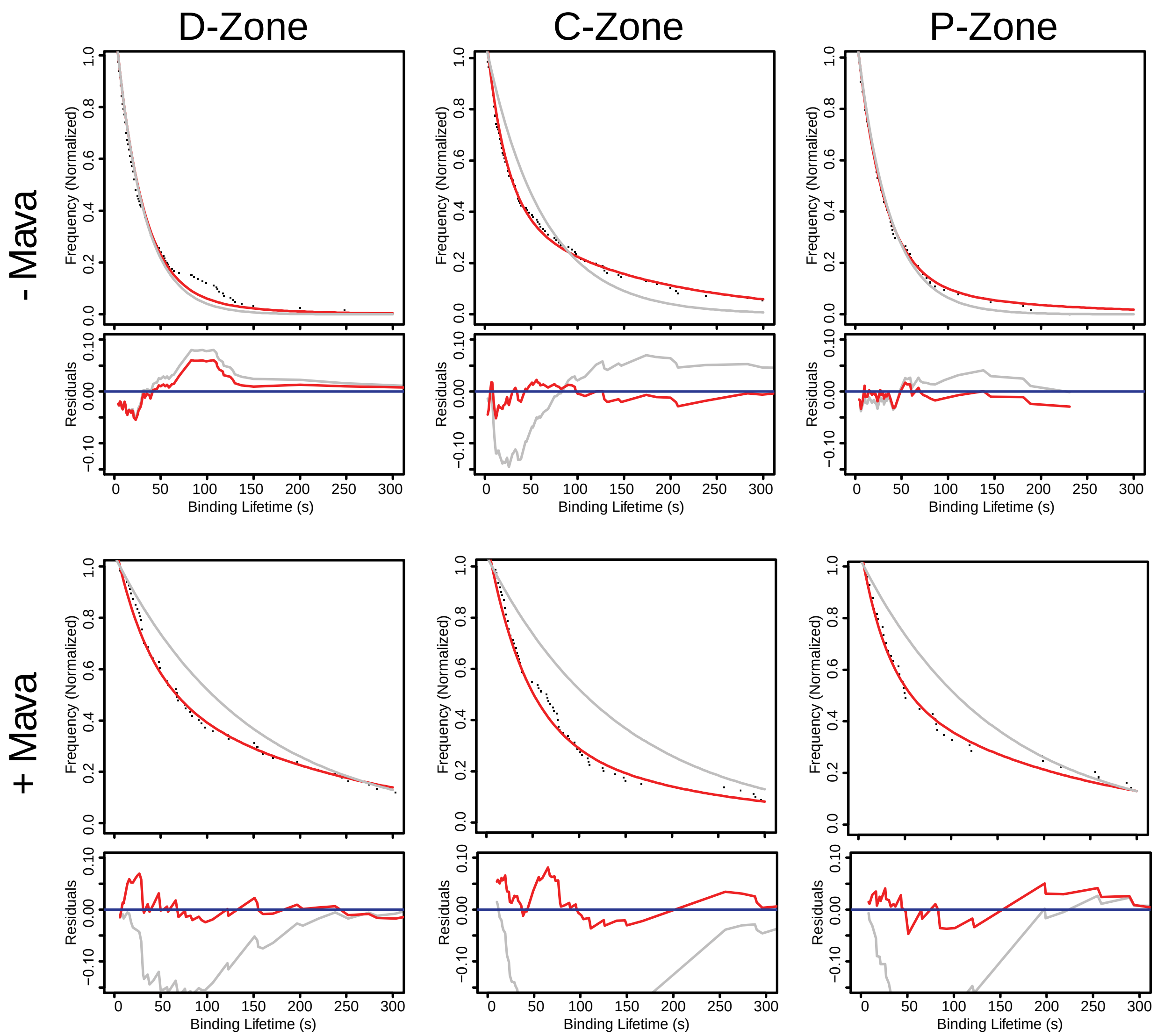


**Figure S4.** Survival plots of D-, C-, and P-zone fluorescent-nucleotide binding event lifetimes with (+Mava) and without Mavacamten (-Mava) as presented in Main Text Figure 3D&E. Lifetimes within each zone are fit with a single (grey line) or double (red lines) exponential function. Residuals are shown below each plot, with matching colors.

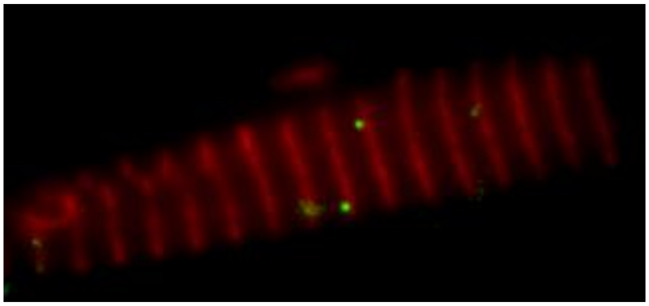

**Movie S1.** Fluorescent ATP binding to myofibril. Rat soleus myofibrils, labeled with anti-myomesin antibodies (red), incubated in Relaxing Buffer (see Materials and Methods). Transient binding of single molecules of BODIPY-ATP (green transient spots) are visible. Video is of 5 minutes of observation, playback is 30X real-time. Field of view measures 32 x 15 microns.
